# Deep Learning-Based Segmentation of 2D Projection-Derived Overlapping Prospore Membrane in Yeast

**DOI:** 10.1101/2025.06.01.656963

**Authors:** Shodai Taguchi, Keita Chagi, Hiroki Kawai, Kenji Irie, Yasuyuki Suda

## Abstract

Quantitative morphological analysis is crucial for understanding cellular processes. While 3D Z-stack imaging offers high-resolution data, the complexity of 3D structures makes direct interpretation and manual annotation challenging and time-consuming, especially for large datasets. Maximum Intensity Projection (MIP) is a common strategy to create more interpretable 2D representations, but this inevitably leads to artificial overlaps between structures, significantly hindering accurate automated segmentation of individual instances by conventional methods or standard deep learning tools. To address this critical challenge in 2D projection analysis, we developed DeMemSeg, a deep learning pipeline based on Mask R-CNN, specifically designed to segment overlapping membrane structures, called prospore membranes (PSMs) during yeast sporulation. DeMemSeg was trained on a custom-annotated dataset, leveraging a systematic image processing workflow. Our optimized model accurately identifies and delineates individual, overlapping PSMs, achieving segmentation performance and derived morphological measurements that are statistically indistinguishable from expert manual annotation. Notably, DeMemSeg successfully generalized to segment PSMs from unseen data acquired from *gip1Δ* mutant cells, capturing the distinct morphological defects in PSMs. DeMemSeg thus provides a robust, automated solution for objective quantitative analysis of complex, overlapping membrane morphologies directly from widely used 2D MIP images, offering a practical tool and adaptable workflow to advance cell biology research.

## Introduction

Understanding the intricate relationship between organelle structure and function is fundamental in cell biology. Microscopy-based morphological analysis offers indispensable insights into diverse cellular processes, encompassing organelle biogenesis, membrane trafficking, cell division, and responses to environmental cues (Nunnari and Suomalainen, 2012; Tojima *et al*., 2024; Voeltz and Prinz, 2007). Quantitative measurements of organelle shape, size, number, and spatial organization are crucial for moving beyond qualitative descriptions towards objective, data-driven conclusions (Peng, 2008; Tojima *et al*., 2024).

While acquiring 3D Z-stack images captures comprehensive spatial information for such detailed analysis, the inherent complexity of these datasets makes manual interpretation and annotation exceptionally challenging and labor-intensive, particularly for large-scale studies (Meijering, 2012). Consequently, projecting 3D data into 2D representations using Maximum Intensity Projection (MIP) is a widely adopted strategy to simplify visualization and analysis.

However, this simplification for interpretability introduces a significant technical hurdle: artificial overlap. Structures distinct in 3D space frequently appear superimposed in the resulting 2D MIP image. This challenge is particularly evident when studying dynamic membrane remodeling processes, such as the formation of prospore membranes (PSMs) during sporulation in budding yeast *Saccharomyces cerevisiae*. PSM formation is a *de novo* process that initiates at meiosis II, driven by the fusion of post-Golgi vesicles at the outer plaque of the spindle pole body (SPB). Subsequently, PSMs undergo a series of morphological changes: they initially appear as distinct dots to form horseshoe-like structures, then elongate into tubules to progressively engulf each of the four haploid daughter nuclei and associated cellular contents. Upon the completion of nuclear engulfment, the leading edge of the membrane fuses and forms spheres, resulting in four spores enclosed within the mother cell cytoplasm (Neiman, 2011, 2024) (**Figure 1A**). The simultaneous growth and shaping of these four membrane structures within the confined cellular space inevitably leads to them being in close proximity. Consequently, when viewed in a 2D MIP image, these distinct PSMs frequently appear to overlap (**Figure 1B**), particularly during the tubular and sphere stages, making it difficult to delineate individual membrane boundaries using traditional automated methods based on thresholding or edge detection (Canny, 1986; Kittler and Illingworth, 1986). This difficulty also complicates manual annotation, which is time-consuming, laborious, and prone to inter- and intra-observer variability, thereby limiting throughput and reproducibility.

**Figure 1.**
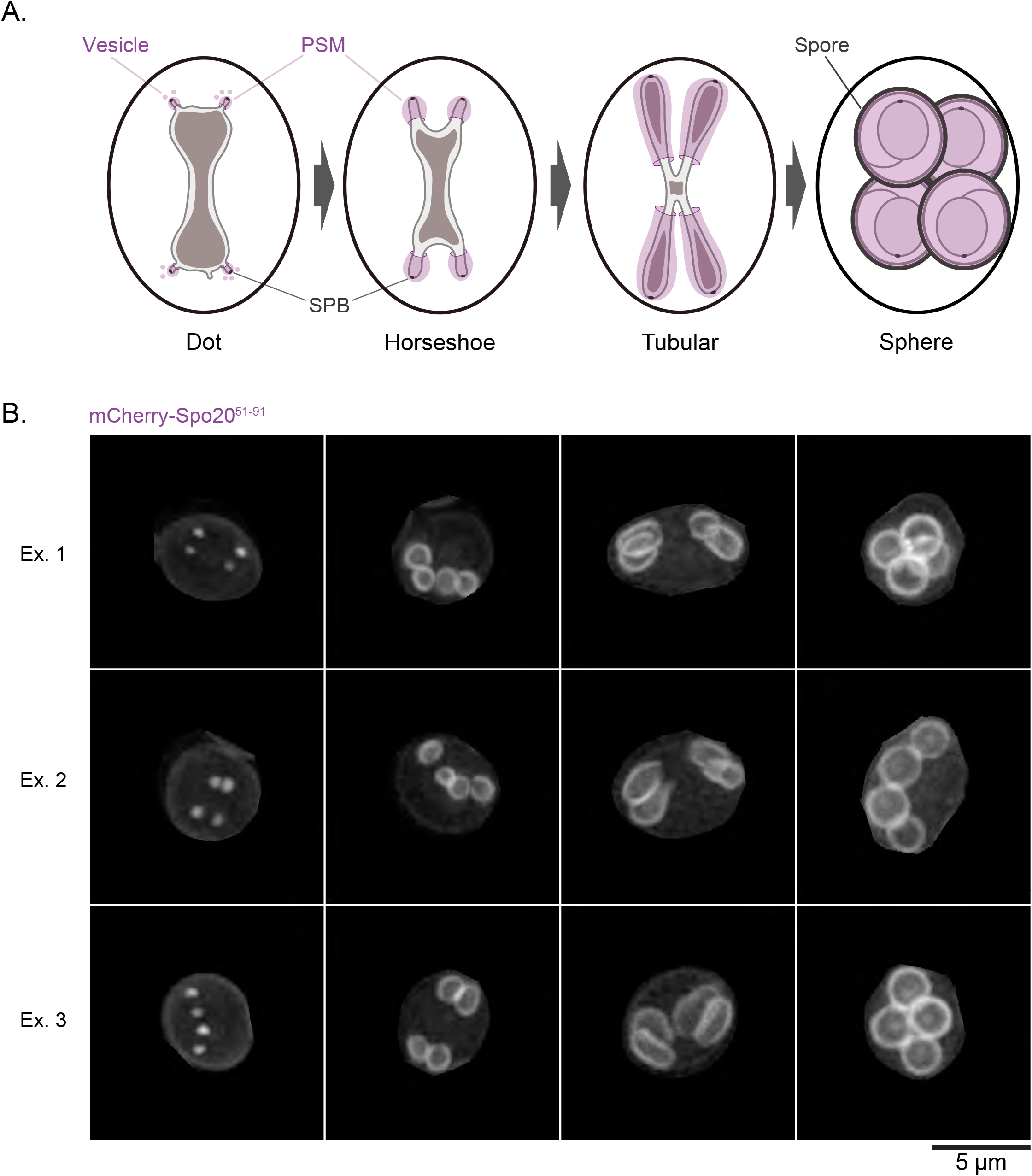
Prospore membrane formation in budding yeast and overlapping 2D image. **(A)** Schematic representation of PSM development during sporulation in *Saccharomyces cerevisiae*. The membranes form and grow to engulf the haploid nuclei, progressing through stages such as Dot, Horseshoe, Tubular, and Sphere, eventually undergoing maturation to form spores. **(B)** Representative 2D MIP fluorescence images of wild-type cells expressing the PSM marker mCherry-Spo20^51–91^. PSM morphologies at each of the stages are shown, illustrating the natural occurrence of overlap between adjacent PSMs in the 2D projection. Scale bar, 5 µm.

The advent of deep learning has revolutionized image segmentation. Seminal architectures like U-Net have demonstrated remarkable success in biomedical image segmentation by effectively capturing contextual information for pixel-level classification (Ronneberger *et al*., 2015). Building on such advances, instance segmentation tools like CellPose (Pachitariu and Stringer, 2022) and ilastik (Berg *et al*., 2019), have greatly improved biological image analysis, they often struggle to precisely segment heavily overlapping subcellular structures in 2D MIP images. This limitation hinders accurate quantitative analysis of individual membrane dynamics such as PSM formation.

To specifically address the segmentation of overlapping structures in 2D MIP images, the Mask R-CNN architecture (He *et al*., 2017) was employed. This architecture is known for its effectiveness in instance segmentation even in complex scenes.

We leveraged the challenging PSM formation process in yeast as an ideal model system to develop and test our approach. The primary objective of this study was to create and validate a Deep learning pipeline based Membrane Segmentation, named DeMemSeg, capable of automatically and accurately segmenting individual, overlapping PSMs from 2D MIP fluorescence images. This involved training the model on a custom dataset generated through meticulous manual annotation of MIP images. Here, we demonstrate that DeMemSeg achieves high segmentation accuracy, comparable to expert manual annotation, and facilitates reliable quantitative analysis of PSM morphology, successfully distinguishing wild-type and mutant phenotypes. This work provides a validated technical approach to overcome the overlap issue inherent in 2D projection analysis, offering a valuable tool and a practical case-study for quantitative cell biology.

## Materials and Methods

### Growth conditions

All yeast strains were cultured at 30°C. Synchronous sporulation was induced as previously described (Neiman, 1998; Suda *et al*., 2024). Briefly, yeast cells freshly grown on synthetic defined (SD, 0.67% Yeast Nitrogen Base without amino acids Difco YNB, 291940; BD Biosciences, Franklin Lakes, NJ, USA) and appropriate amino acids, 2% dextrose (D(+)-glucose, 045-31167; FUJIFILM Wako Pure Chemical Corporation, Tokyo, Japan)) plates containing 2% agar (STAR Agar L-grade 01, RSU-AL01-500G; Rikaken Co., Tokyo, Japan) were inoculated into 3 mL SD liquid medium and grown overnight. 0.7 mL of overnight culture was transferred to 15 mL YPA (2% Peptone (Bacto Peptone, 211677; Thermo Fisher Scientific, Waltham, Massachusetts, US), 1% Yeast Extract (Bacto Yeast Extract, 212750; Thermo Fisher Scientific), and 0.003% Adenine (6-Aminopurine, 012-11512; FUJIFILM Wako Pure Chemical Corporation, Tokyo, Japan), 2% potassium acetate (167–3185; FUJIFILM Wako Pure Chemical Corporation) liquid medium and grown for 16 hours. Cells were harvested, washed once in distilled water, then resuspended in sporulation medium (2% potassium acetate) at OD_600_ = 2.0, and incubated with vigorous shaking. After six hours, *NDT80* expression was induced by adding 2 mM β-estradiol (E8875; Sigma-Aldrich, Massachusetts, US) diluted in ethanol to a final concentration of 2 μM.

### Yeast strains and plasmids

Standard genetic manipulation techniques were employed unless stated otherwise. All yeast strains utilized in this study are derived from the SK1 strain background. Strains and plasmid used are listed in Supplementary Table 1. The following alleles and plasmids were constructed in previous studies: *NDT80::hphNT1::P*_*4xlexA*_*-9xMyc-NDT80 AUR1::P*_*ACT1*_-LexA-ER - haVP16::AUR1-C (Nakamura *et al*., 2021; Tachikawa *et al*., 2001).

### Fluorescence microscopy and image analysis

Static fluorescence images of sporulating yeast cells expressing the PSM marker were acquired using a Leica Thunder Imager Live Cell system equipped with a 100x oil immersion objective (NA 1.4) and an sCMOS camera (DFC9000 GTC, Leica Microsystems). Yeast cells were induced for sporulation and imaged approximately every 30 minutes to capture various stages of PSM development. Yeast cultures were spotted onto a 24 × 50 mm cover slip (No.1, Cat. No. C024501; Matsunami Glass Ind., Ltd., Osaka, Japan) and sandwiched with a smaller 18 × 18 mm cover slip (No.1, Cat. No. C218181; Matsunami Glass Ind., Ltd.). For each field of view, Z-stacks consisting of 40-50 slices were acquired at the space of 0.21 μm intervals. These 3D Z-stack images were subsequently subjected to the image restoration using the Leica Thunder Imager software, followed by Maximum Intensity Projection (MIP) to generate 2D images (**Figure 2A**). These 2D MIP images were used for the construction of the training dataset and for subsequent image analysis (**Figure 2B, C**). For the model generality test, additional images were also acquired using a Leica TCS SP8 LIGHTNING confocal microscope and a Keyence BZ-X710 all-in-one fluorescence microscope. Images acquired from these microscopes were not used in the training stage.

**Figure 2.**
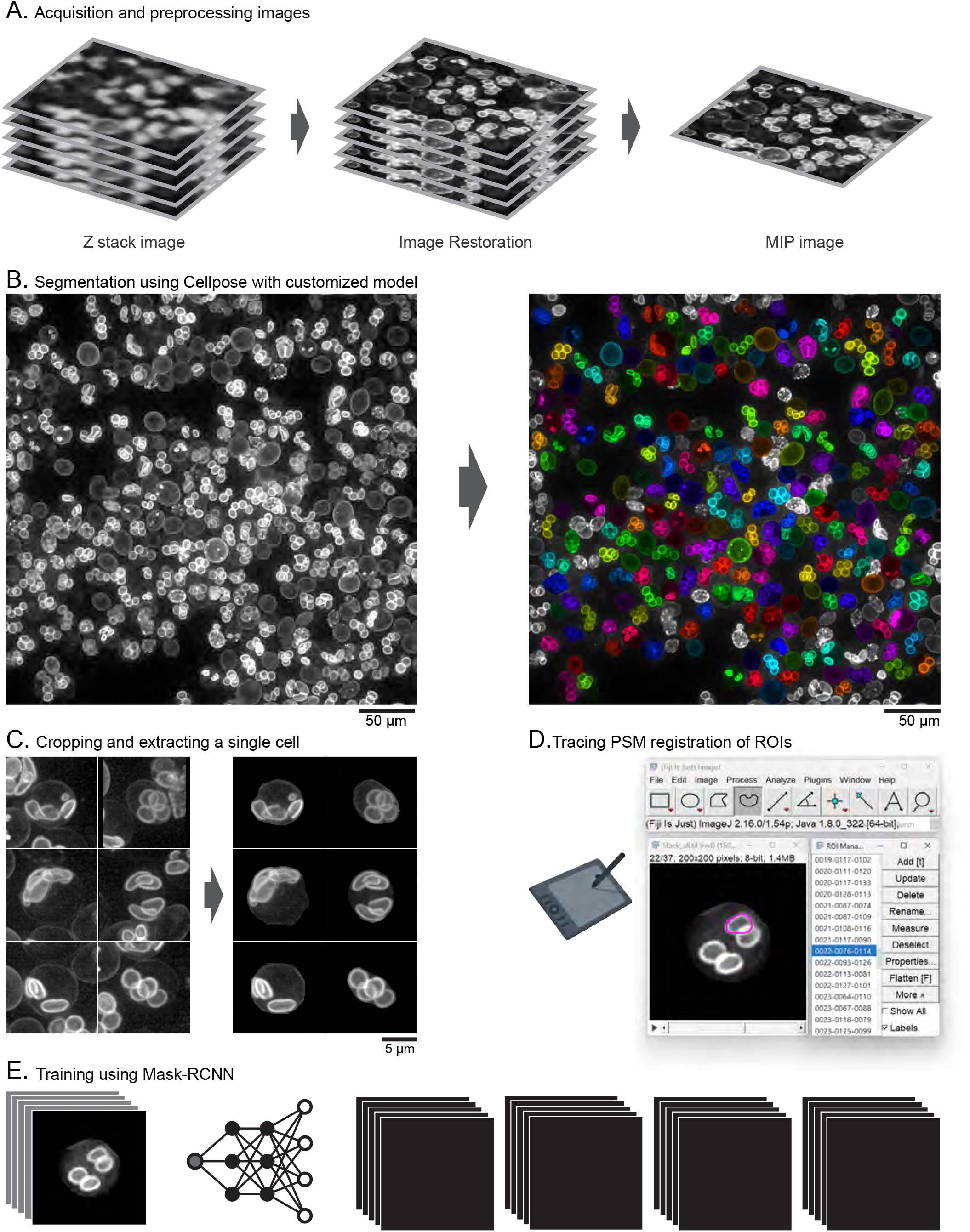
Pipeline of DeMemSeg and workflow for construction of the annotated training dataset. **(A)** Image preprocessing workflow. Raw 3D Z-stack images acquired from fluorescence microscopy are subjected to image restoration, followed by Maximum Intensity Projection (MIP) to produce a 2D image. **(B)** Whole-cell segmentation using a custom-trained CellPose 2.0 model for the isolation of individual cells. **(C)** Cropping of single-cell regions from the MIP images, guided by CellPose segmentation masks. **(D)** Manual tracing of individual PSM contours within cropped single-cell images utilizing ImageJ/Fiji. **(E)** Generation of image-mask pairs. An original cropped single-cell image (left) is processed by a deep neural network to produce corresponding binary masks for each PSM instance. These pairs constitute the training data.

### DeMemSeg Training Dataset Construction

An annotated dataset was generated for training the DeMemSeg model. Crucially, all images used for creating this training, validation, and testing dataset were derived from wild-type SK1 strain yeast cells; images from the *gip1*Δ mutant strain were not included in any part of this dataset preparation or model training process. The overall workflow is depicted in **Figure 2**.

#### Acquisition and preprocessing images (Figure 2A)

As described previously, 2D MIP images were generated from 3D Z-stacks after image restoration. An example of such a wide-field MIP image, serving as the input for the subsequent processing steps.

#### Segmentation using Cellpose with customized model (Figure 2B)

To facilitate the annotation of individual PSMs within single cells, MIP images were first processed using a custom-trained CellPose 2.0 model (Pachitariu and Stringer, 2022). This model was specifically trained to segment whole yeast cells, identifying valid, complete cells while excluding those partially cut off by the image border or identified as dead cells. This step helped to isolate analyzable cells from dense fields.

#### Single-Cell Image Cropping (Figure 2C)

Based on the masks from the custom CellPose model, 200 x 200 pixel regions centered on the centroid of each valid cell were cropped from the original MIP images. The corresponding CellPose mask was used to blank out signals from any neighboring cells or background within the crop, ensuring that each cropped image predominantly featured a single cell. This was done to simplify the subsequent manual annotation task and potentially reduce false positives during model inference.

#### Manual Annotation of Prospore Membranes (Figure 2D)

We manually traced the precise contours of individual PSMs within these cropped single-cell images. This was performed using ImageJ/Fiji software (Schindelin *et al*., 2012) running on a PC, with an iPad Air 5th generation (Apple Inc., CA, US) connected via remote desktop software (spacedesk, Version 2.1.43) to serve as a drawing tablet interface for precise tracing. Each traced PSM boundary was saved as an ImageJ/Fiji Region of Interest (ROI). Even when PSMs overlapped in the 2D projection, each was delineated as a distinct ROI.

#### Dataset Finalization and Splitting (Figure 2E)

The saved ROIs were converted into binary mask images. The final dataset comprised pairs of an original cropped single-cell MIP image and its corresponding PSM mask image(s) (a single cell image could have up to four PSM masks). This collection of images and instance masks was then formatted into the COCO dataset structure (Lin *et al*., 2014) to be compatible with the MMdetection framework. This curated dataset, consisting of 7040 ROIs across all PSM developmental stages, was used for model training. The dataset was split into training (80%), validation (10%), and testing (10%) sets, ensuring unbiased evaluation of the model’s performance on unseen data.

### DeMemSeg Model Architecture, Training, and Inference

#### Model Architecture

DeMemSeg is based on the Mask R-CNN architecture (He *et al*., 2017), a well-established framework for instance segmentation. The model was implemented using the MMdetection toolbox (Chen *et al*., 2019). We primarily used a ResNet-50 and a ResNet-101 backbone (He *et al*., 2015), pre-trained on the ImageNet dataset (Deng *et al*., 2009).

#### Training Parameters and Evaluation Metrics

Standard data augmentation, specifically random horizontal flipping (applied with 50% probability), was applied to the training data to enhance model robustness and generalization. The model was trained using a Stochastic Gradient Descent (SGD) optimizer (Ruder, 2016) with an initial learning rate of 0.0025 and with batch sizes of 4 and 8. Training typically proceeded for up to 24 epochs. Model performance was monitored on the validation set using standard COCO evaluation metrics for instance segmentation. The primary metric was segm_mAP, defined as the mean Average Precision (mAP) calculated over a range of Intersection over Union (IoU) thresholds from 0.5 to 0.95 with a step of 0.05 (often denoted as mAP@[.5:.05:.95]). We also specifically report segm_mAP_50 (mAP at an IoU threshold of 0.5) and segm_mAP_75 (mAP at an IoU threshold of 0.75) to provide further insight into performance at different levels of localization accuracy.

#### Inference Parameters

For model inference, during post-processing, the IoU threshold for Non-Maximum Suppression (NMS) was set to 0.8. The maximum number of detected instances per image was limited to 4, reflecting the biological context of up to four PSMs per sporulating yeast cell. The prediction score threshold for accepting an instance as a valid detection was optimized on the validation dataset and subsequently applied when evaluating the test dataset.

### SAM 2 Model for Comparison

For qualitative comparison of segmentation performance on time-lapse images, we utilized SAM 2 (Segment Anything Model 2) (Ravi *et al*., 2024). Given SAM 2’s recognized capabilities in robust video segmentation and its potential utility for tracking cellular structures over time, we considered it a relevant state-of-the-art model for qualitative comparison against DeMemSeg’s performance on our time-lapse PSM data. Briefly, for the time-lapse sequences, positive point prompts were manually provided to SAM 2 for each individual PSM visible in a reference frame, typically at the horseshoe stage. These prompts were then propagated or applied across all slices (frames) of the time-lapse sequence. Inference was performed both forwards and backwards in time from the initially prompted frame to generate segmentations for the entire sequence. Default parameters for the SAM 2 model were used unless otherwise specified.

### Morphological Parameter Quantification and Statistical Analysis

Morphological parameters, specifically PSM length (perimeter of the segmented mask) and roundness (calculated as 4×π×Area/Perimeter^2^), were extracted from both manual annotations and DeMemSeg predictions using custom scripts in Python and Fiji/ImageJ. Statistical significance of differences between groups for these parameters was assessed using Welch’s t-tests. P-value annotations are as follows: ns (not significant) p > 0.05; *: 0.01 < p ≤ 0.05; **: 0.001 < p ≤ 0.01; ***: 0.0001 < p ≤ 0.001; ****: p ≤ 0.0001.

To directly assess the concordance between manual and automated measurements on a per-object basis, a correlation analysis was also performed. For each cell, individual PSM instances from the manual annotation set were paired with corresponding instances from the DeMemSeg prediction set by identifying the pair with the maximum Intersection over Union (IoU). These paired measurements for PSM length were then visualized using scatter plots, and the strength of the linear correlation was quantified by calculating the coefficient of determination (R^2^). Furthermore, to provide a qualitative view of segmentation performance, 50 representative instance pairs were selected and visualized. These examples were chosen by ranking all pairs by their IoU and selecting the top 10 (best), bottom 10 (worst), and 30 randomly sampled instances from the remainder.

### Computation resource

All model training and prediction tasks for both DeMemSeg (using CellPose and MMdetection) and SAM 2 were performed on a workstation equipped with an NVIDIA GeForce RTX 3060 Ti GPU (8GB VRAM).

### Code availability

The Python scripts, Jupyter notebooks, custom model weights (for DeMemSeg and the CellPose pre-processing model), and example data used in this research are available from the GitHub repository: https://github.com/MolCellBiol-tsukuba/DeMemSeg.

## Results

### Image Processing Pipeline for Training DeMemSeg and Analyzing Prospore Membranes

Accurate quantitative analysis of dynamic membrane structures like prospore membranes (PSMs) from 2D MIP images (**Figure 1B**) requires precise instance segmentation, often preceded by a standardized image processing and dataset preparation workflow. To facilitate both the training of our DeMemSeg model and subsequent PSM analysis, we established a pipeline that can be applied for similar quantitative analyses of membrane morphology.

The initial step involves processing wide-field MIP images containing numerous cells (**Figure 2A**) using a custom-trained CellPose 2.0 model to segment individual yeast cells (**Figure 2B**). This whole-cell segmentation serves two key purposes: first, it allows for the filtering of unsuitable cells (e.g., those at image borders or dead cells, as detailed in Materials and Methods); second, it provides the basis for isolating individual cells for focused analysis. Subsequently, 200 x 200 pixel regions centered on the centroid of each valid cell are cropped. Critically, to ensure that DeMemSeg’s training and inference are focused solely on the PSMs within a single target cell, the corresponding CellPose mask is applied to blank out signals from any neighboring cells or background within each crop (**Figure 2C**). For training dataset generation, PSMs within single-cell cropped images were manually annotated using ImageJ/Fiji, with each PSM instance defined as a distinct Region of Interest (ROI) (**Figure 2D**). These ROIs were converted into binary masks, forming image-mask pairs that were structured into a COCO format dataset for training the Mask R-CNN based DeMemSeg model (**Figure 2E**). The detailed procedures for image acquisition, CellPose model training, manual annotation, data augmentation, and dataset splitting are provided in Materials and Methods. This systematic approach to data preparation ensures consistent, high-quality input for both training the segmentation model and its application in quantitative analysis.

### DeMemSeg Model Optimization and Initial Performance Validation

To identify a suitable minimum dataset size for robust performance and determine the optimal balance between model complexity, computational resources, and segmentation accuracy for the DeMemSeg model, we systematically evaluated the impact of training dataset size and key architectural hyperparameters. All evaluations utilized the segm_mAP (COCO mAP@[.5:.05:.95]) metric on the validation set.

First, to understand the relationship between data quantity and model performance, we investigated the effect of training dataset size. Using a ResNet-50 backbone and a batch size of 4, we trained DeMemSeg with varying percentages of the total available training data, from 2% (141 ROIs) up to 100% (7040 ROIs) (**Figure 3A**). As expected, segm_mAP generally improved with increasing data. Notably, even with 50% of the data (3520 ROIs), the model achieved approximately 0.662, a performance level close to that obtained with 100% of the data (approx. 0.670). This suggests that a dataset of around 3500 ROIs can yield substantial performance for this task. Furthermore, the learning curves for both 50% and 100% dataset sizes began to plateau around 12-15 epochs. Next, we explored the influence of batch size and backbone architecture, using 50% of the training data (3520 ROIs) for these comparisons (**Figure 3B**). Batch size in deep learning refers to the number of training examples utilized in one iteration. Comparing ResNet-50 with batch size 4 (R50 bs4) against batch size 8 (R50 bs8), we observed that R50 bs4 (segm_mAP ≈ 0.662) slightly outperformed or performed comparably to R50 bs8 (segm_mAP ≈ 0.650), while being more memory efficient.

**Figure 3.**
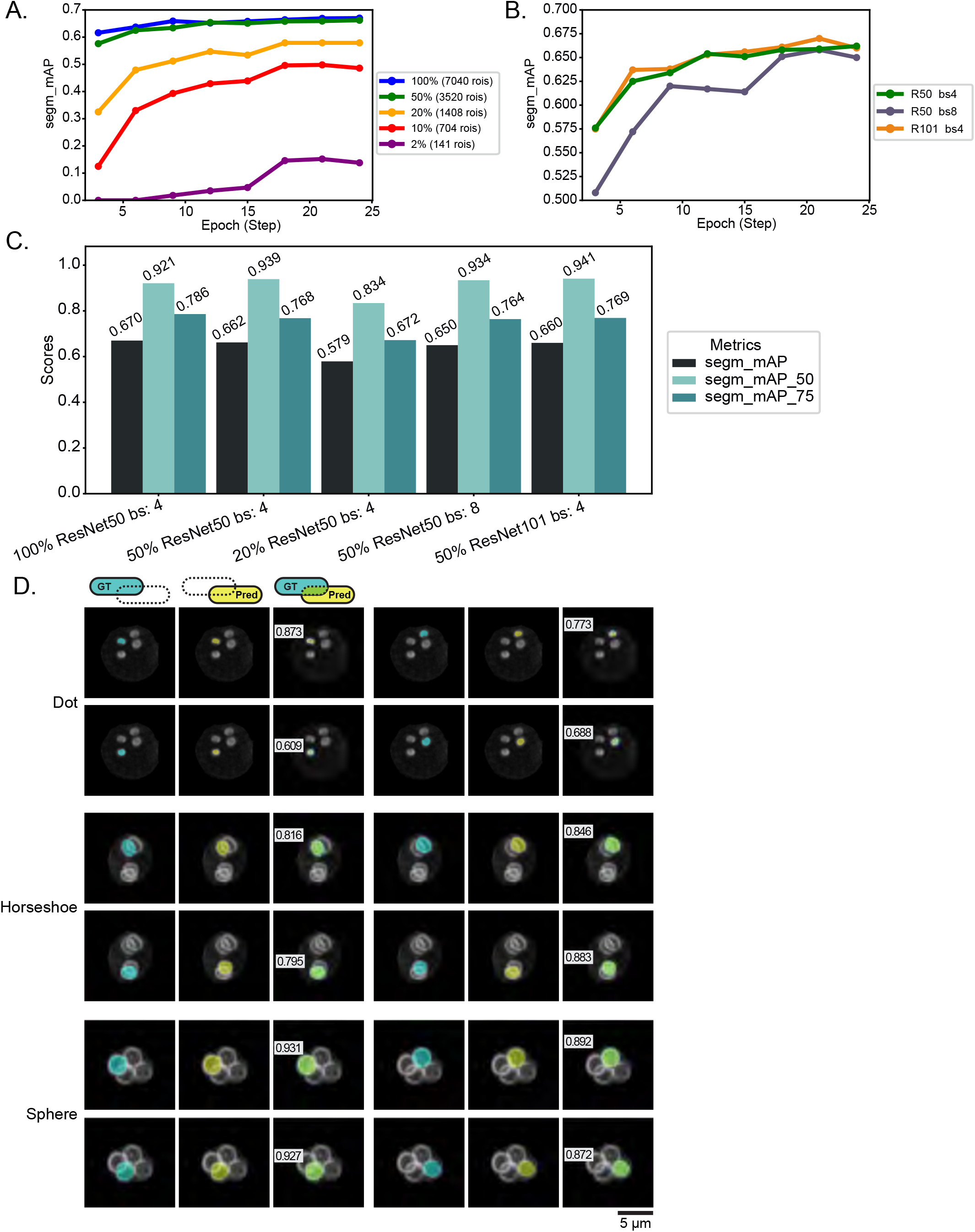
Optimization of DeMemSeg model parameters and visual validation of segmentation accuracy. **(A)** Impact of training dataset size on model performance. The graph plots segm_mAP (COCO mAP@[.5:.05:.95]) on the validation set against training epochs. Each colored line represents a model trained with a different percentage of the total training data. All models utilized a ResNet-50 backbone and a batch size of 4. **(B)** Impact of model configuration (backbone and batch size) on performance. The graph plots segm_mAP (COCO mAP@[.5:.05:.95]) on the validation set against training epochs. Each colored line represents a different model configuration (ResNet-50 with batch size 4 [R50 bs4], ResNet-50 with batch size 8 [R50 bs8], or ResNet-101 with batch size 4 [R101 bs4]). All models were trained using 50% of data. **(C)** Bar graph comparing key segmentation performance metrics (segm_mAP, segm_mAP_50 [mAP at IoU 0.50], segm_mAP_75 [mAP at IoU 0.75]) for different DeMemSeg model configurations. Configurations shown are: 100% training data with ResNet-50 backbone and batch size 4 (100% ResNet50 bs:4); and models trained with 50% or 20% of data using ResNet-50 bs:4, or with 50% data using ResNet-50 bs:8 or ResNet-101 bs:4. **(D)** Representative examples of DeMemSeg’s segmentation performance on test images for Dot, Horseshoe, and Sphere stages of PSM development. Each instance is shown as a triplet: the GT mask (cyan), DeMemSeg’s prediction mask (Pred, yellow), and a combined view of both GT and Pred, on the MIP image. Intersection over Union (IoU) score is also shown. Scale bar, 5 µm.

We then compared different backbone architectures: ResNet-50 and ResNet-101 (**Figure 3B**). ResNet architectures are distinguished by their depth; ResNet-101 is a deeper network with more layers and consequently more parameters and a higher theoretical representational capacity compared to ResNet-50, but it also demands greater computational resources, particularly GPU memory. Due to GPU memory limitations imposed by the increased parameter size of ResNet-101, this backbone was evaluated only with a batch size of 4 (R101 bs4). Comparing the performance of ResNet-50 with batch size 4 (R50 bs4, segm_mAP ≈ 0.662) with that of ResNet-101 with batch size 4 (R101 bs4, segm_mAP ≈ 0.660), we found no significant performance improvement with the deeper ResNet-101 backbone for this specific task and dataset size.

A summary of key performance metrics (segm_mAP, segm_mAP_50, and segm_mAP_75) for these critical configurations is presented in **Figure 3C**. While the 50% ResNet101 bs:4 configuration achieved the highest segm_mAP_50 (0.941), our primary goal of precise PSM shape analysis necessitates high accuracy at stricter IoU thresholds. Therefore, we focused on segm_mAP_75 as a critical indicator of precise boundary delineation. The configuration using 100% of the training data with a ResNet-50 backbone and batch size 4 yielded the highest segm_mAP_75 score of 0.786, outperforming other configurations (e.g., 50% ResNet101 bs:4 achieved segm_mAP_75 of 0.769). Consequently, to ensure the highest possible accuracy for subsequent detailed morphological analyses, we selected the model trained on 100% of the data with the ResNet-50 backbone and a batch size of 4 as the final DeMemSeg model. This configuration also provided the highest overall segm_mAP (0.670).

Following optimization and parameter finalization, segmentation with the selected DeMemSeg model yielded prediction masks from representative test images. These prediction masks were compared to manually created ground truth (GT) masks to verify segmentation accuracy, as shown in **Figure 3D**. Four distinct prediction instances for PSM are depicted at each stage of PSM development (Dot, Horseshoe, Tubular, and Sphere). Visual inspection across PSM development reveals almost complete overlap between DeMemSeg’s prediction masks and the GT masks and achieved relatively high IoU scores (0.609 to 0.873 in Dot stage, 0.795 to 0.883 in Horseshoe stage, and 0.872 to 0.931 in Sphere stage). These robust mAP scores and high individual IoU scores suggested that the optimized DeMemSeg model accurately segments individual PSMs with high fidelity for detailed quantitative analysis.

### Visual Validation of DeMemSeg’s Segmentation on Time-Lapse Image and Comparison with SAM 2

To further assess DeMemSeg’s robustness and its generalization capability to unseen images, the optimized model was applied to time-lapse MIP image sequences acquired from sporulating yeast cells in which continuous progression of PSM proceeds. For the comparison, SAM 2 (Ravi *et al*., 2024), a prominent foundation model, was also applied to the same image sequences. As depicted in **Figure 4A**, a consistent temporal framework was employed to compare the outputs from the original image sequence, SAM 2 predictions (**Supplementary Video 1**), and DeMemSeg predictions (**Supplementary Video 2**). This comparison spanned a series of 13 time points at 5-minute intervals, covering a total duration of 0 to 120 minutes.

**Figure 4.**
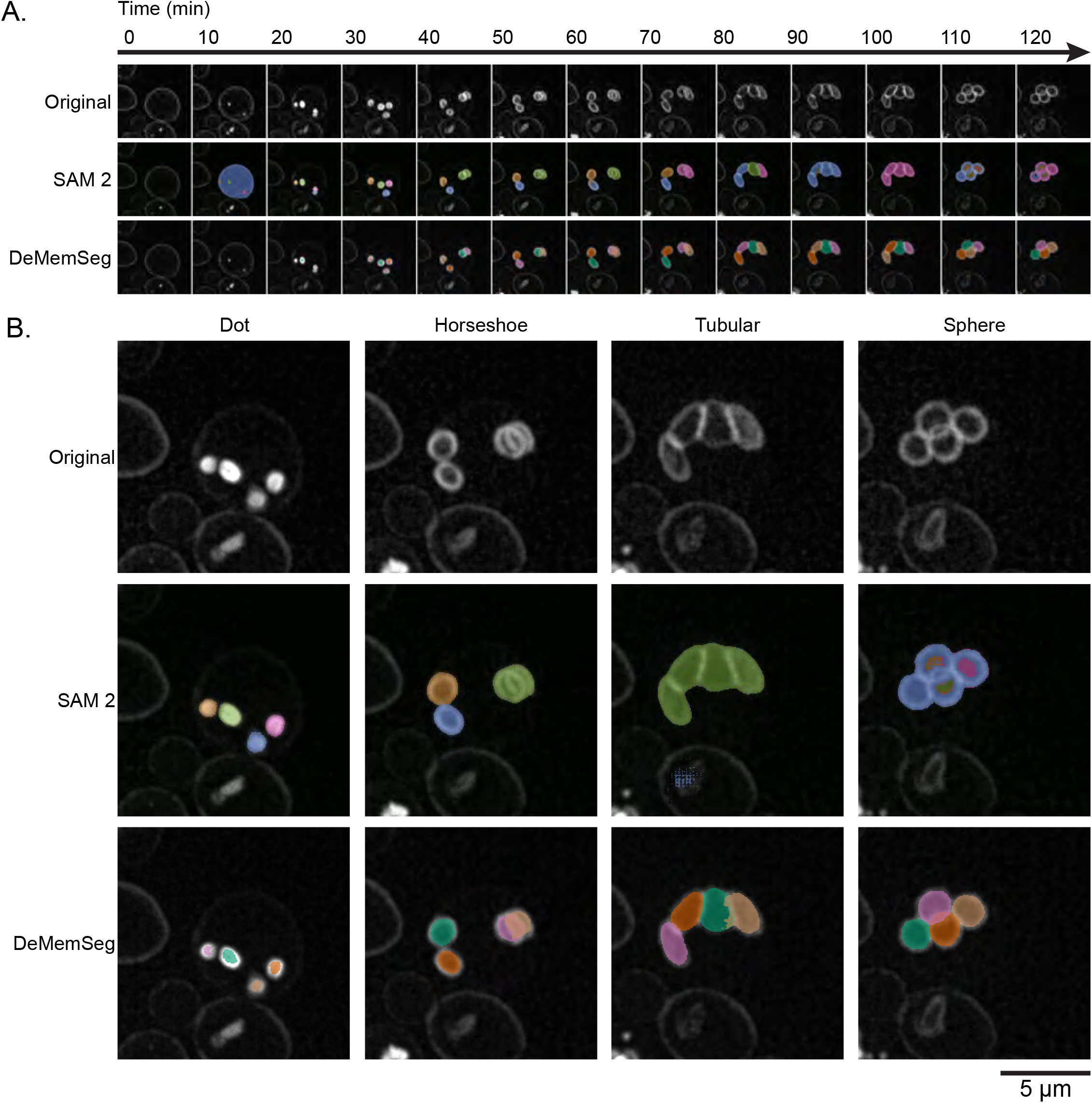
Qualitative evaluation of DeMemSeg on untrained time-lapse image snapshots. **(A)** Time-lapse fluorescence image displaying the temporal progression of PSM development (0-120 minutes) in WT cell expressing mCherry-Spo20◻^1^◻◻^1^ as observed in the original image sequence, SAM 2 predictions (see **Supplementary Video 1**), and DeMemSeg predictions (see **Supplementary Video 2**). **(B)** Representative MIP snapshot images selected from (A), showing distinct PSM developmental stages: Dot, Horseshoe, Tubular, and Sphere. The images are not used for the training of the model. Scale bar, 5 µm.

Representative snapshots of key morphological stages of PSM formation: Dot, Horseshoe, Tubular, and Sphere selected from these time-lapse image sequences are presented in **Figure 4B**. As a foundation model, SAM 2 is designed for broad applicability and can often perform zero-shot segmentation on objects for which it was not explicitly trained. Indeed, SAM 2 was able to identify the general regions corresponding to PSMs, even though PSMs were likely not part of its original training corpus. However, SAM 2 frequently encountered difficulties in precisely separating adjacent or overlapping PSMs into distinct instances (**Figure 4A, 4B, SAM 2, Supplementary Video 1**). On the other hand, DeMemSeg successfully identified and delineated individual PSMs across all stages of PSM formation, and maintained the ability to separate instances even when PSMs existed in close proximity or exhibited significant overlap (**Figure 4A, 4B, DeMemSeg, Supplementary Video 2**).

### DeMemSeg Enables Accurate Quantitative Phenotyping of Wild-Type and Mutant PSMs

Having established DeMemSeg’s foundational accuracy and its robustness on untrained dynamic sequences, we next assessed its utility for detailed quantitative morphological analysis in the mutant defective in PSM formation. Deletion of *GIP1*, coding for a meiosis-specific targeting subunit of type 1 protein phosphatase (PP1), exhibits the defects in PSM elongation and spore maturation (Nakamura *et al*., 2017; Suda *et al*., 2024; Tachikawa *et al*., 2001). Images from *gip1*Δ cells were not included in the training dataset, allowing evaluation of DeMemSeg’s generalization.

Visual inspection of DeMemSeg’s segmentation outputs confirmed its ability to accurately delineate PSMs across various developmental stages not only in WT cells but also in the morphologically distinct *gip1*Δ mutant cells (**Figure 5A**). The model successfully identified and segmented smaller and more irregularly shaped PSMs that are found in the *gip1*Δ cells. To quantitatively substantiate this observation, we performed a per-object correlation analysis of PSM length between manual and DeMemSeg-derived measurements. This analysis confirmed a strong linear correlation for both WT (R^2^ = 0.715) and *gip1Δ* (R^2^ = 0.699) strains, providing direct evidence of the model’s accuracy on an individual instance level (**Figure S1**). Representative images of these 50 segmented instances are provided in **Figure S2** for WT and **Figure S3** for *gip1Δ* cells.

**Figure 5.**
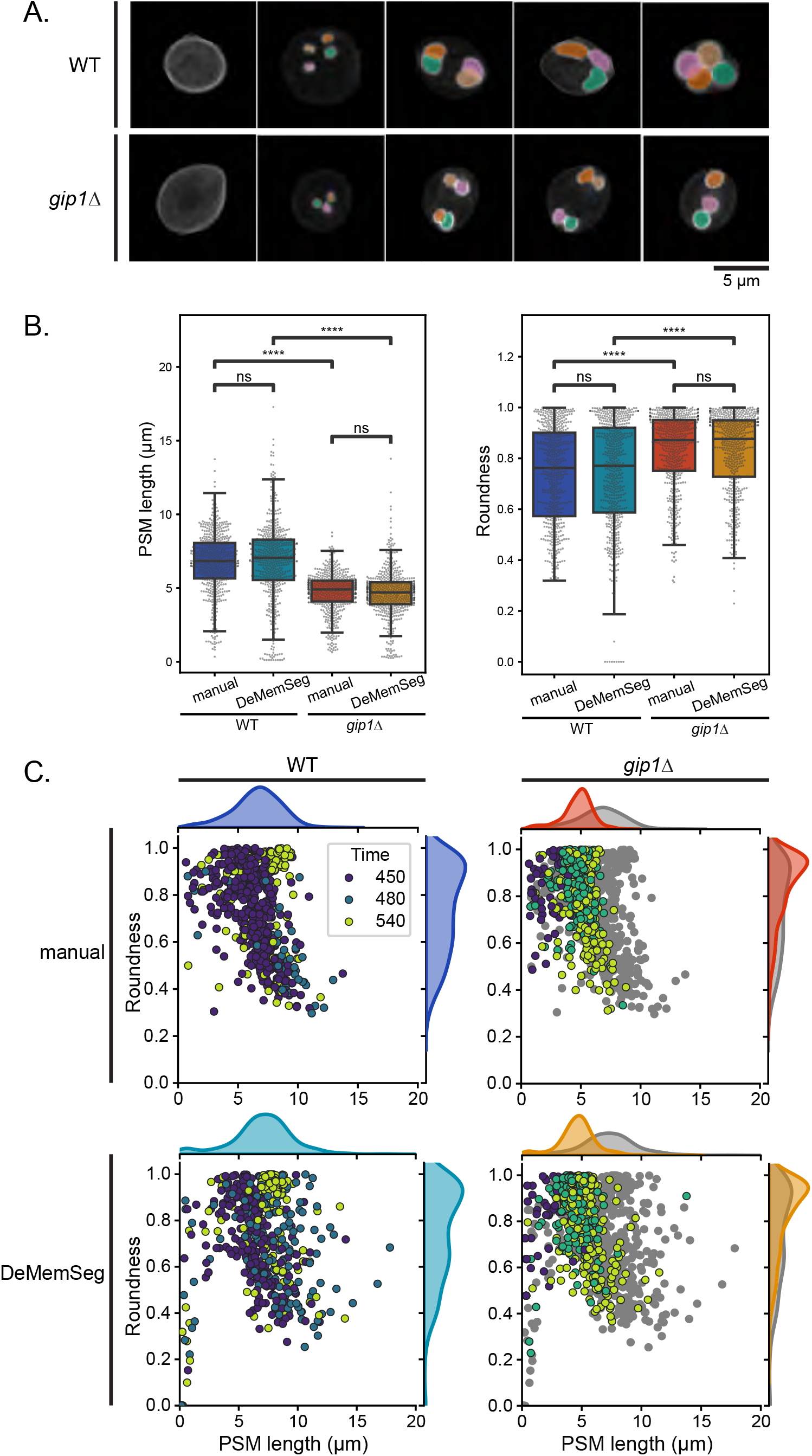
DeMemSeg enables accurate and detailed morphological characterization of wild-type and *gip1*Δ PSM. **(A)** Representative MIP images of original and corresponding DeMemSeg segmentation predictions for wild-type (WT) and the *gip1Δ* mutant cells expressing mCherry-Spo20◻^1^◻◻^1^. Scale bar, 5 µm. **(B)** Box plots showing the comparisons of PSM length (µm, left panel) and roundness (dimensionless, right panel) for WT and *gip1Δ* strains. Each point within the box plots represents a measurement from an individual PSM. Statistical significance (Welch’s t-test) is indicated: ns, p > 0.05; *, 0.01 < p ≤ 0.05; ****, p ≤ 0.0001. **(C)** Scatter plots visualizing PSM length versus roundness, with marginal density plots for each parameter. Top and bottom row panels show manual annotations and DeMemSeg predictions, respectively. For *gip1Δ* panels, WT data from WT is also presented as gray points for comparison. Data points are colored according to the image acquisition time (Time (min)) as indicated by the color legend.

Quantitative comparisons of key morphological parameters—PSM length (perimeter) and roundness, where each data point represents a single PSM—were performed using both manual annotations and DeMemSeg predictions for WT and *gip1*Δ cells (**Figure 5B**). For both WT and *gip1*Δ cells, measurements for PSM length and roundness showed no statistically significant difference (ns) between manual annotations and DeMemSeg predictions (PSM length: WT p = 0.4627, *gip1*Δ p = 0.4866; PSM roundness: WT p = 0.7077, *gip1*Δ p = 0.6723). This demonstrates DeMemSeg’s reliability in extracting these features with human-level accuracy for both normal and aberrant morphologies. Moreover, DeMemSeg’s automated predictions effectively captured the known morphological phenotype of the *gip1*Δ mutant. As shown in **Figure 5B**, PSMs in *gip1Δ* were significantly shorter (mean of PSM length predicted by DeMemSeg: 4.65 µm for *gip1Δ* vs. 6.61 µm for WT; p = 2.437e-52, ****) and significantly more round (mean of roundness predicted by DeMemSeg 0.83 for *gip1*Δ vs. 0.72 for WT;: p = 1.028e-16, ****) than those in WT when comparing DeMemSeg’s predictions for both strains. This finding is consistent with the comparison of manual annotations (mean of PSM length: 4.72 µm for *gip1Δ* vs. 6.64 µm for WT; p = 4.239e-71, ****; mean of roundness: 0.83 for *gip1Δ* vs. 0.72 for WT; p = 1.555e-24, ****), highlighting DeMemSeg’s capability to accurately quantify these distinct biological phenotypes.

Further insights into the morphological distinctions and temporal progression were obtained from 2D scatter plots of PSM length versus roundness, combined with marginal density plots for each parameter, where data points are colored by image acquisition time (**Figure 5C**). Distributions derived from manual annotations are presented in the top row of panels, while the bottom row shows those from DeMemSeg predictions. These scatter plots effectively visualize a characteristic trajectory in the dynamic progression of PSM morphology over time. In WT cells after 450 min, most of PSMs showed sphere-like dots with short perimeters and slightly elongated horseshoes. Subsequently, at 480 min, PSMs reach their maximum perimeter and exhibit their lowest roundness values, reflecting their extended cup-like structures. By 540 min, a significant population of WT PSMs is concentrated in the region characterized by a PSM length of approximately 10 µm, indicating the formation of spherical, mature spores in the Sphere stage.

In contrast, the *gip1*Δ mutant exhibits a markedly different temporal and morphological trajectory. While some progression in PSM length is observed, the mutant PSMs largely fail to reach the same elongation state observed in WT. Even at 540 minutes, PSMs of *gip1*Δ cells remain less elongated and exhibit diverse roundness (including almost 1.0). These observations recapitulate our previous findings that *gip1*Δ cells exhibit defects in PSMs elongation but achieve normal membrane closure, resulting in small, spherical, immature spores (Nakamura *et al*., 2017; Park and Neiman, 2012; Suda *et al*., 2024).

Importantly, these characteristic phenotypic alterations observed in the *gip1Δ* mutant were consistently represented in both the manual annotation data (top row) and the DeMemSeg prediction data (bottom row) (**Figure 5C**). This comprehensive analysis, incorporating temporal progression, demonstrates that DeMemSeg not only accurately segments individual PSMs but also enables quantitative characterization of complex morphological phenotypes, providing a powerful scheme for advancing our understanding of cellular processes and the impact of genetic perturbations.

## Discussions

In this study, we developed DeMemSeg, a Mask R-CNN-based deep learning pipeline, to accurately segment overlapping prospore membranes (PSMs) from 2D MIP images, a significant challenge in quantitative cell biology. Systematic optimization yielded a robust model with high concordance with ground truth annotations and strong IoU scores for individual instances.

A key contribution of DeMemSeg is its ability to facilitate precise quantitative analysis of complex biological phenotypes. The DeMemSeg’s morphological measurements were statistically indistinguishable from expert manual annotations for both wild-type (WT) and the *gip1Δ* mutant which is not included in the training data, underscoring its reliability and generalization capability. DeMemSeg effectively captured and quantified the known morphological defects in *gip1Δ* cells (Nakamura *et al*., 2017; Park and Neiman, 2012; Suda *et al*., 2024), highlighting its utility for objective phenotypic characterization. Furthermore, our qualitative comparisons on untrained time-lapse data bring to light an important consideration regarding specialized versus generalist models. While foundation models like SAM 2 exhibit impressive broad generalization, tasks involving unique challenges with specific biological structures—such as the artificial overlaps created in 2D MIP images from 3D Z-stacks—can significantly benefit from custom-trained models. DeMemSeg, having been specifically trained on our PSM dataset which includes numerous examples of such MIP-induced overlaps, developed a more nuanced understanding of these particular boundary conditions. This suggests that for achieving high-fidelity segmentation in specialized scenarios, a customized model like DeMemSeg can offer more effective performance than a general-purpose foundation model.

The automated segmentation of challenging structures by DeMemSeg significantly enhances efficiency, objectivity, and potential for large-scale morphological analyses, offering substantial improvements over manual methods. Specifically, this automation substantially reduces the time expenditures associated with laborious manual segmentation, human judgment errors, and inter-observer variability. This contributes to the production of highly consistent and reproducible datasets. Ultimately, these advancements in laboratory automation lower the barrier to addressing large-scale microscopy datasets. This enables researchers to address more complex biological questions that were previously intractable due to analytical limitations. This work offers not only a validated tool for researchers studying yeast meiosis and sporulation but also a practical case study for those investigating membrane dynamics and organelle morphology in other cellular contexts.

Despite these achievements, certain limitations are considered. As a supervised model, DeMemSeg’s optimal performance is tied to its training data, and application to significantly different contexts may require retraining or fine-tuning. Our analysis of the pre-trained model on images from different microscopy systems (**Figure S4**) suggests that while the performance is highest on images with clear object boundaries (such as those enhanced by image restoration), it retains a degree of useful functionality on images from other commercial microscopy systems, provided the PSM outlines are reasonably visible. It is also important to note that while the pre-trained model has some generalizability, optimal performance on images from a new system may require retraining the model on a custom-annotated dataset using the provided pipeline. The plateau observed in learning curves also suggests that exploring more diverse data augmentation techniques could further enhance generalization.

Moreover, while 2D MIP analysis is practical for visualizing the overall morphology of structures, it inherently loses 3D spatial information. Therefore, caution is necessary when inferring precise 3D spatial relationships between different structures using segmentation masks derived from 2D MIP images. Apparent co-localization or containment in the 2D projection may not accurately reflect the true 3D arrangement.

In conclusion, DeMemSeg provides a powerful and validated automated solution for segmenting and quantifying overlapping membrane structures in widely used 2D MIP images. Its demonstrated capacity to accurately characterize complex phenotypes facilitates a deeper, more quantitative understanding of cellular organization and dynamics, offering a valuable asset for advancing image-based research in cell biology.

## Abbreviations

2D: Two-Dimensional
3D: Three-Dimensional
AI: Artificial Intelligence
AP: Average Precision
mAP: mean Average Precision
bs: Batch Size
CNN: Convolutional Neural Network
DL: Deep Learning
IoU: Intersection over Union
LED: Light-Emitting Diode
MIP: Maximum Intensity Projection
ML: Machine Learning
NA: Numerical Aperture
PSM: Prospore Membrane
SPB: Spindle Pole Body
R50: ResNet-50
R101: ResNet-101
ROI: Region of Interest
SAM 2: Segment Anything Model 2
sCMOS: scientific Complementary Metal-Oxide-Semiconductor
SGD: Stochastic Gradient Descent
WT: Wild-Type
OD_600_: Optical Density at 600 nm

## Acknowledgments

We would like to express our sincere gratitude to Haruka Ozaki, Yoshinori Hayakawa and lab members of bioinformatics laboratory in University of Tsukuba, lab members of the Laboratory of Molecular Cell Biology at University of Tsukuba, and Hiroyuki Tachikawa (Rikkyo University) for insightful discussions and valuable advice throughout the course of this research.

## Data Availability Statement

The Python scripts, Jupyter notebooks, and model weights that were used in the research and the data analyses are available from the GitHub repository: https://github.com/MolCellBiol-tsukuba/DeMemSeg

## Author Contribution Statement

Conceptualization: ST; Data curation: ST; Formal Analysis: ST; Funding acquisition: YS; Investigation: ST; Methodology: ST, KC, HK, YS; Project Administration: ST, YS; Resources: ST, KI, Y.S; Software: ST; Supervision: KC, HK, YS; Validation: ST; Visualization: ST; Writing – Original Draft: ST, KC, HK, KI, YS; Writing – Review & Editing: ST, KC, HK, KI, YS All authors have read and agreed to the published version of the manuscript.

## Supplementary Video Legend

**Supplementary Video 1:** This video shows the application of the SAM 2 model to a time-lapse image sequence of a sporulating yeast cell expressing mCherry-Spo20◻^1^◻◻^1^ shown in Figure 4A. SAM 2’s segmentation predictions are overlaid on the original fluorescence images. The sequence corresponds to the time points indicated at the top right, and a scale bar is shown at the bottom right.

**Supplementary Video 2:** This video shows the application of the trained DeMemSeg model to a time-lapse image sequence of a sporulating yeast cell expressing mCherry-Spo20◻^1^◻◻^1^ shown in Figure 4A. DeMemSeg’s segmentation predictions are overlaid on the original fluorescence images, demonstrating its ability to segment individual, overlapping PSMs throughout their dynamic development from Dot to Sphere stages. The sequence corresponds to the time points indicated at the top right, and a scale bar is shown at the bottom right.

