## Supplementary Figure for "Deep Learning-Based Segmentation of 2D Projection-Derived Overlapping Prospore Membrane in Yeast"

Figure S1

A. Scatter plot for IoU max PSM pairs in WT

B. Scatter plot for IoU max PSM pairs in *gip1*Δ

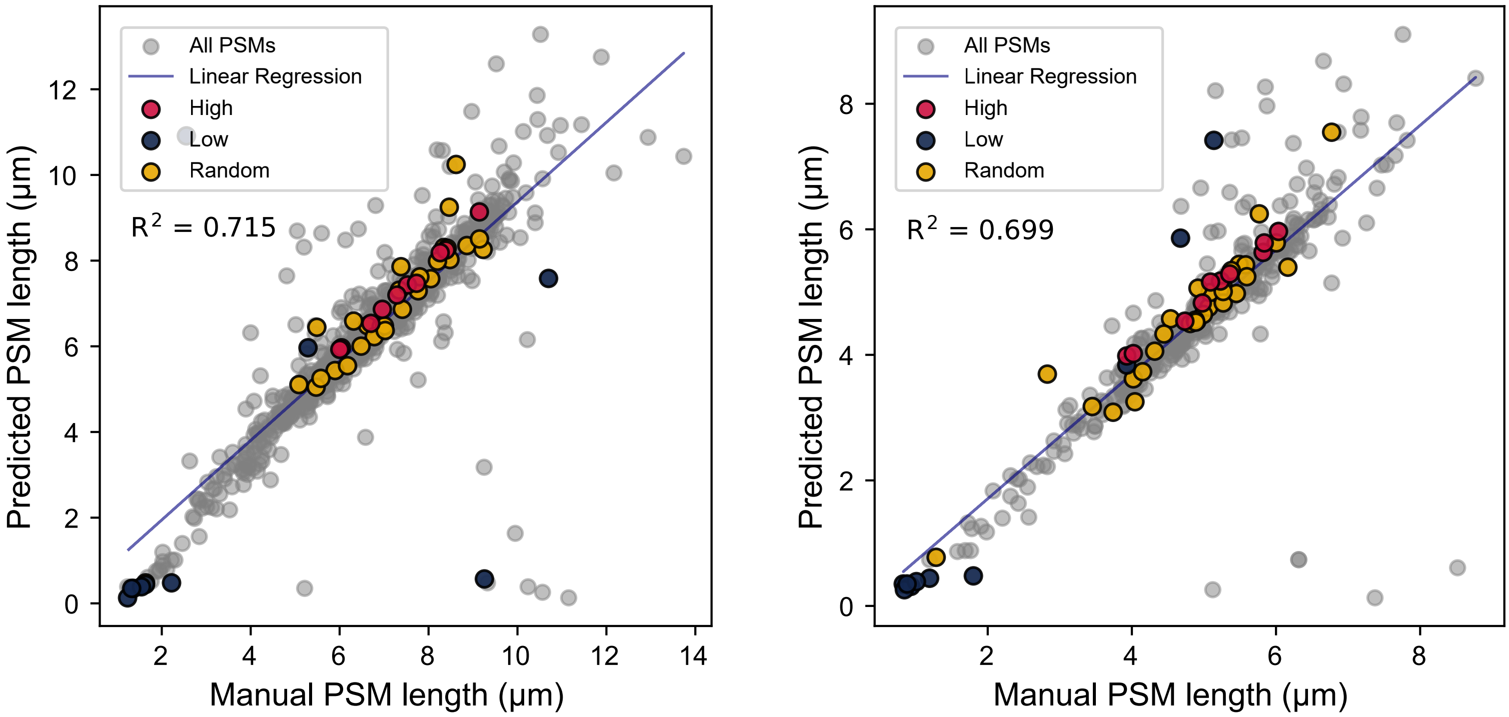


**Supplementary Figure Legends**

**Figure S1. Correlation analysis of PSM length between manual and DeMemSeg measurements.** Scatter plots comparing the prospore membrane (PSM) length measured from manual annotations (x-axis) and DeMemSeg predictions (y-axis) for (A) wild-type (WT) and (B) *gip1*Δ strains. Each point represents a single PSM instance, paired based on the maximum Intersection over Union (IoU) between manual and predicted masks within the same cell. Points highlighted in red, yellow, and navy blue correspond to the high IoU (top 10 pairs), low IoU (worst 10 pairs), and randomly sampled PSM pairs visualized in Figure S2 and S3, respectively. The blue line represents the linear regression fit, with its corresponding $R^2$ value displayed below the label.

**Figure S2. Visual examples of DeMemSeg segmentation performance on wild-type (WT) cells.** Representative images of 50 individual prospore membranes (PSMs) from WT cells, comparing manual annotations (cyan), DeMemSeg predictions (yellow), and their merge (light green). The examples were selected based on the Intersection over Union (IoU) ranking. The figure displays: (A) the 10 instances with the highest IoU, (B) the 10 instances with the lowest IoU, and (C) 30 instances randomly sampled from the remainder. For each instance, the IoU value and the PSM length (in µm) from both manual and DeMemSeg measurements are shown.

**Figure S3. Visual examples of DeMemSeg segmentation performance on *gip1*Δ mutant cells.** Representative images of 50 individual prospore membranes (PSMs) from *gip1*Δ cells, comparing manual annotations (cyan), DeMemSeg predictions (yellow) and their merge (light green). The examples were selected based on the Intersection over Union (IoU) ranking. The figure displays: (A) the 10 instances with the highest IoU, (B) the 10 instances with the lowest IoU, and (C) 30 instances randomly sampled from the remainder. For each instance, the IoU value and the PSM length (in µm) from both manual and DeMemSeg measurements are shown.

Figure S2

A. WT PSMs with the top 10 highest IoU values

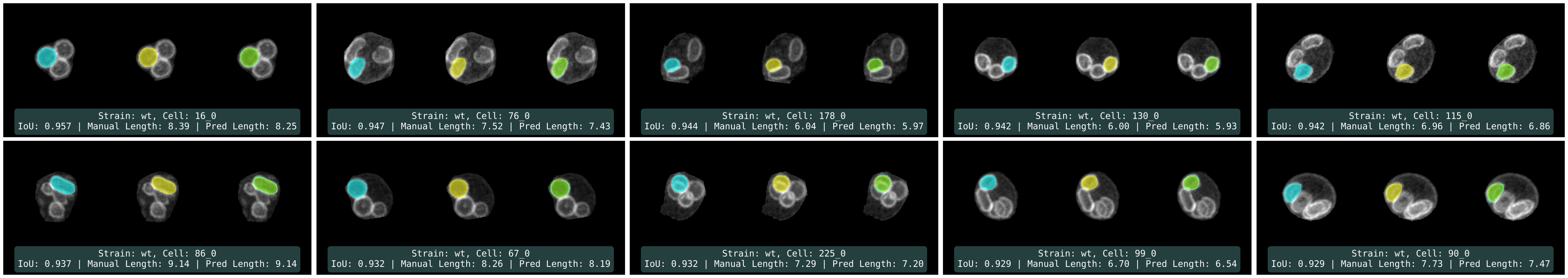


B. WT PSMs with the 10 lowest IoU values

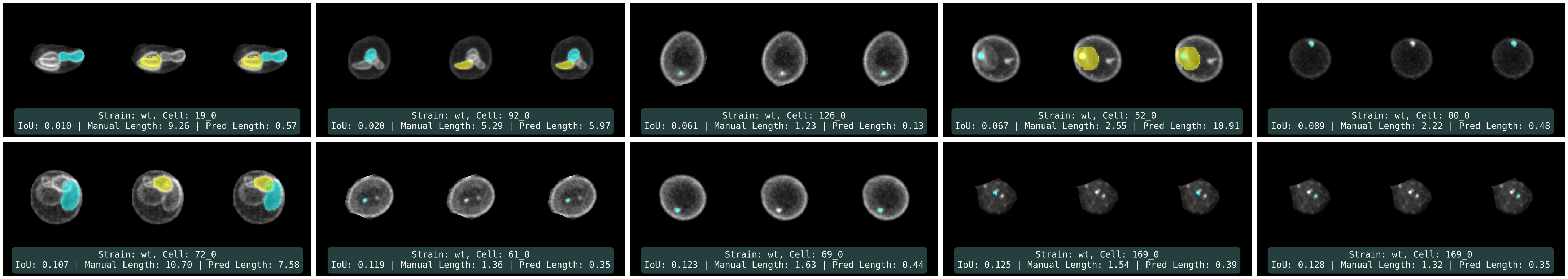


C. WT PSMs randomly sampled (30 samples)

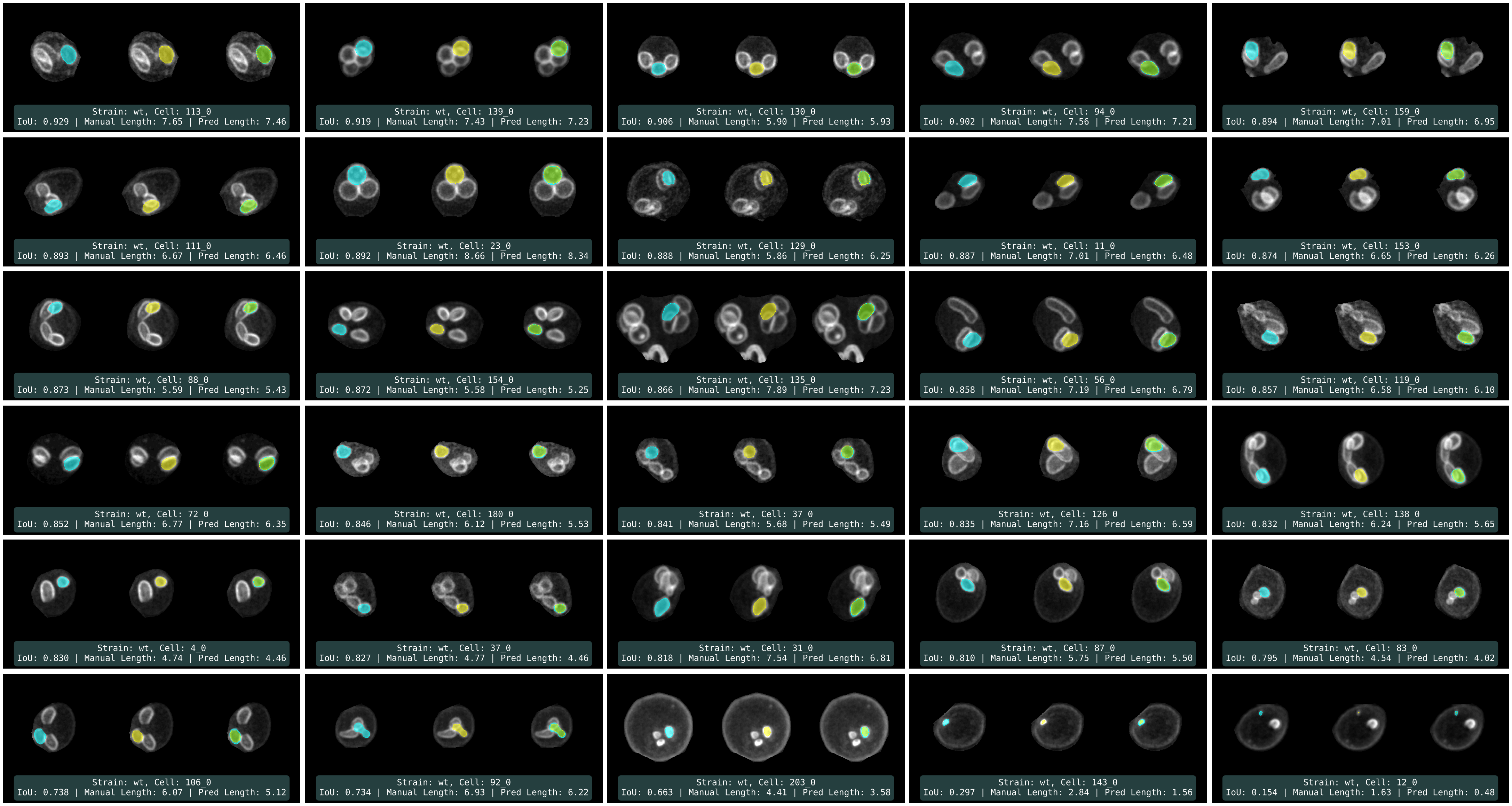

Figure S3

A. *gip1Δ* PSMs with the top 10 highest IoU values

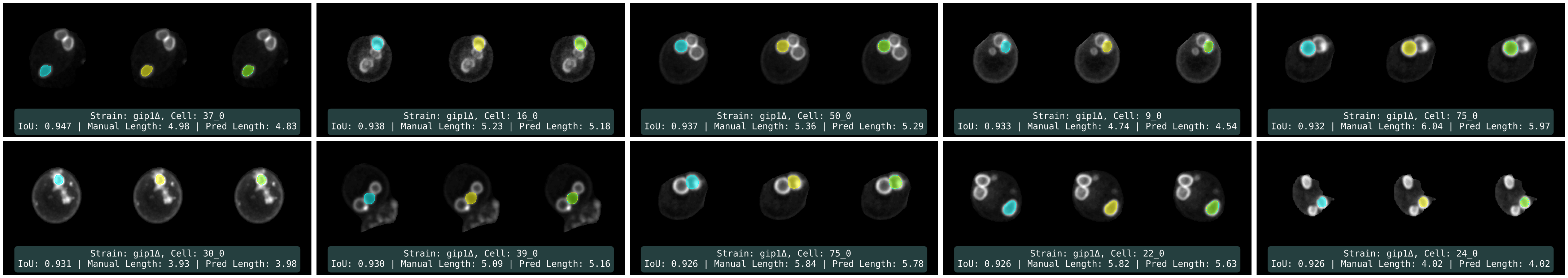


B. *gip1Δ* PSMs with the 10 lowest IoU values

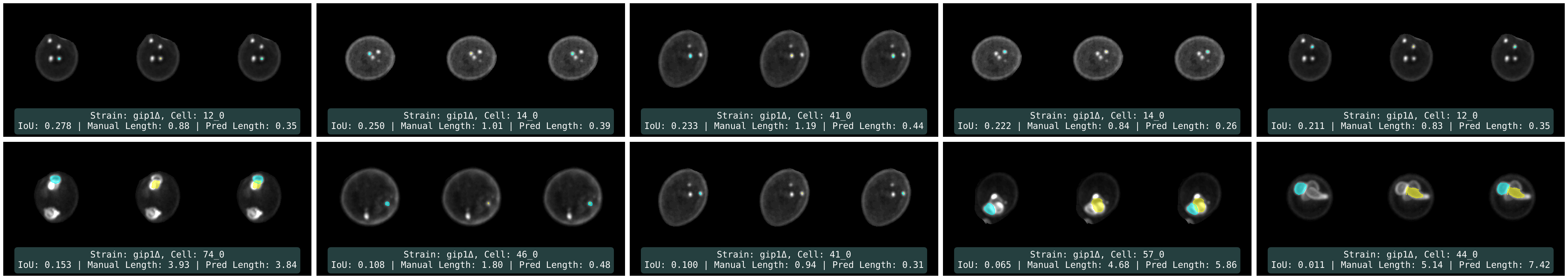


C. *gip1Δ* PSMs randomly sampled (30 samples)

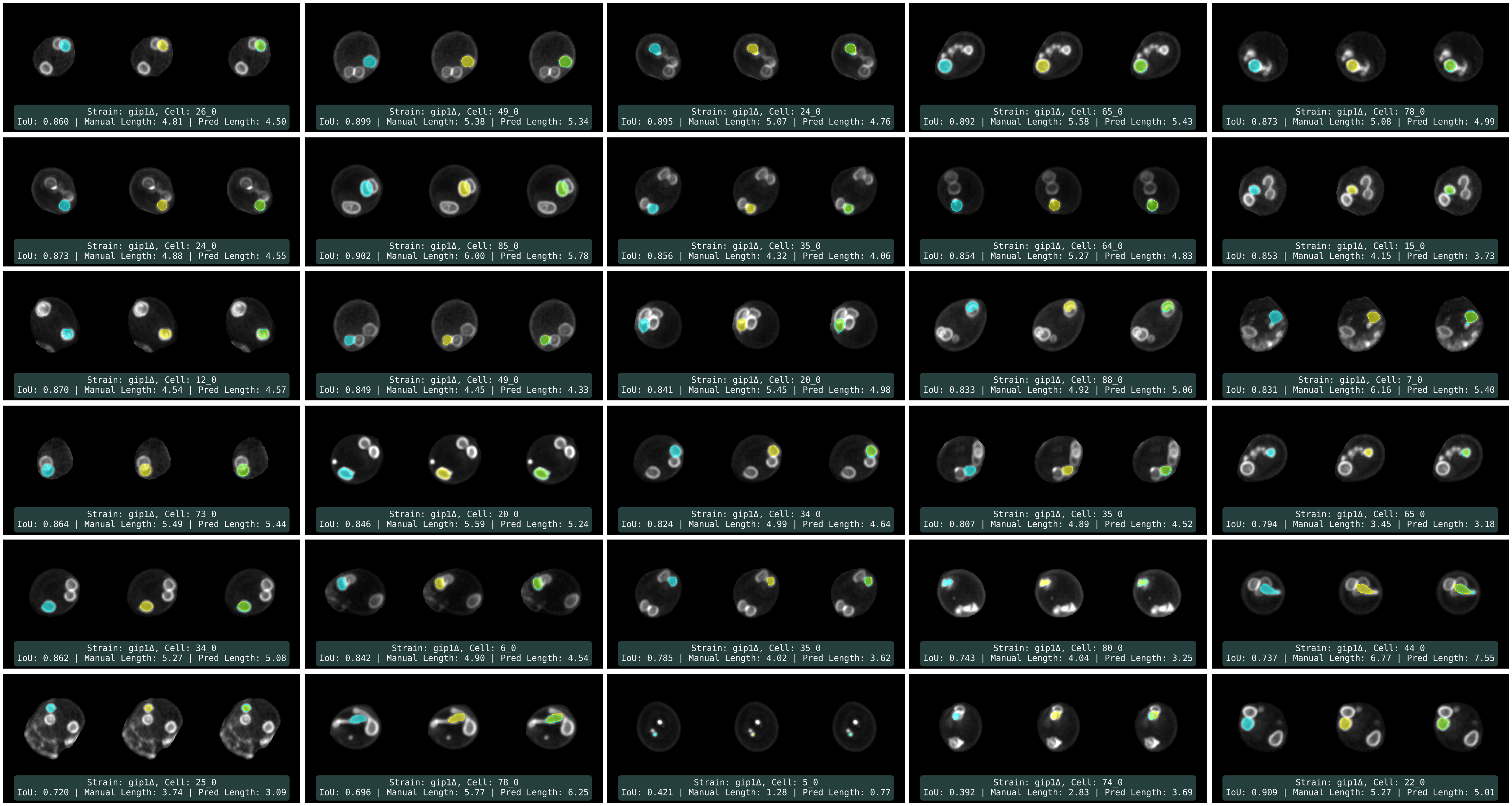

Figure S4

A. Leica THUNDER Imageing Systems, without THUNDER (Image Restoration)

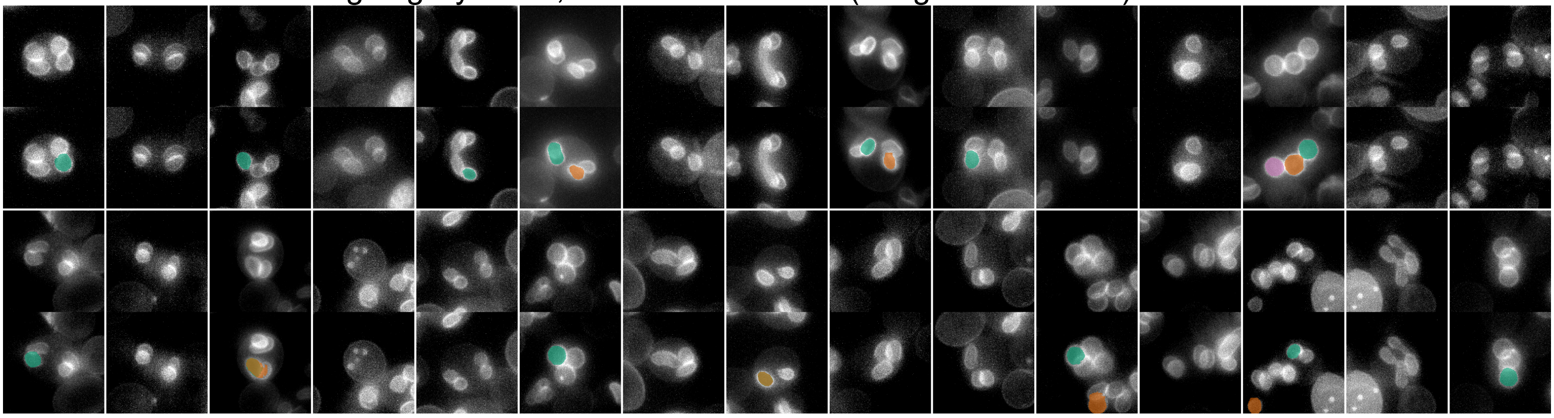

B. Leica SP8 LIGHTNING Confocal Microscope

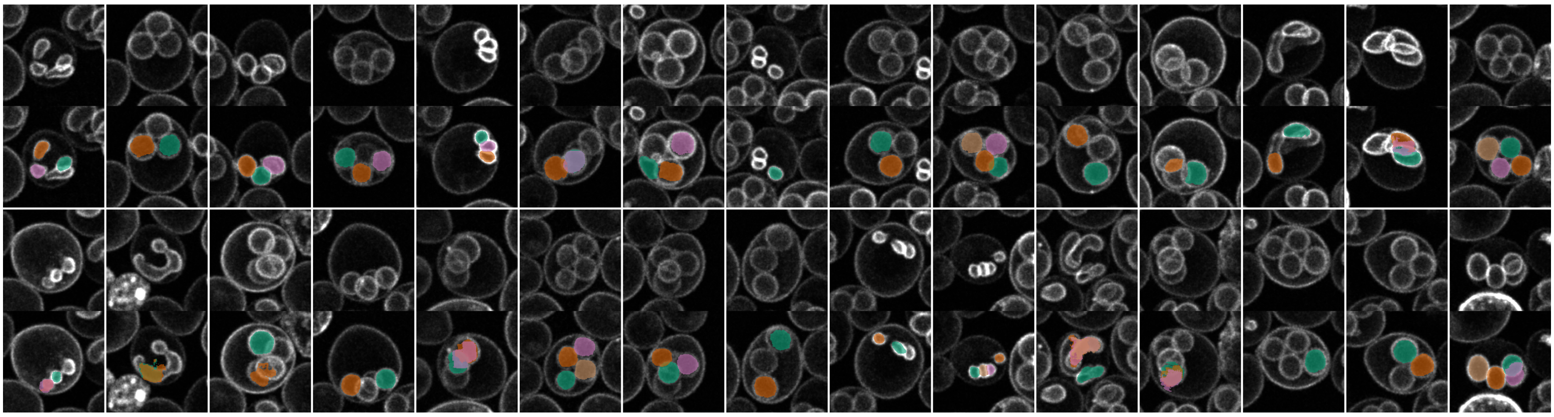

C. Keyence All-in-One Fluorescence Microscope - BZ-X710

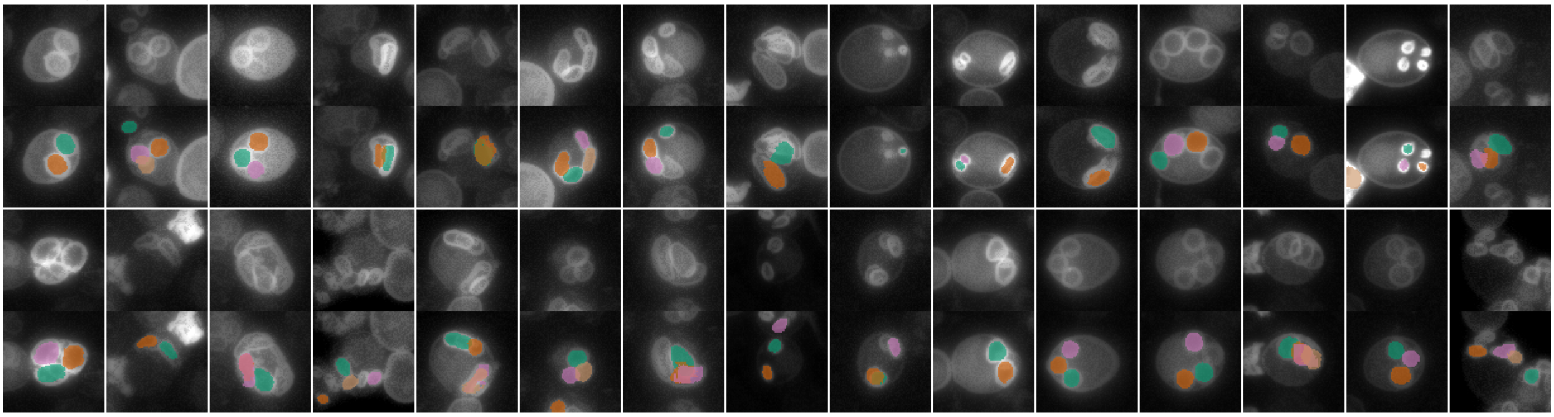

**Figure S4. Performance of the pre-trained DeMemSeg model on images from different microscopy systems and conditions.**

Representative images showing the application of the pre-trained DeMemSeg model to test images acquired from different microscopy systems and settings. The model was not retrained for these examples. In each panel, the top rows show the original images, and the bottom rows show the corresponding segmentation outputs. (A) Images from a Leica THUNDER system without image restoration. (B) Images from a Leica SP8 LIGHTNING confocal microscope. (C) Images from a Keyence BZ-X710 all-in-one fluorescence microscope.
