## Supplementary Table 1 for "Deep Learning-Based Segmentation of 2D Projection-Derived Overlapping Prospore Membrane in Yeast"

**Supplementary table 1. Yeast strains and plasmids used in this study.**

Strain Name	Genotype	Source

TNY1218	*MAT***a**/*MAT*α *his3*Δ*SK*/*his3*Δ*SK ura3*/*ura3::pRS306-mCherry-Spo20^51-91^ trp1::hisG*/*trp1::hisG leu2*/*leu2 arg4-NspI/ARG4*

*lys2/lys2 ho*Δ*::LYS2/ho*Δ*::LYS2 rme1::LEU2/RME1*

*AUR1::P_ACT1_-LexA-ER-haVP16::AUR1-C/AUR1::P_ACT1_-LexA-ER-haVP16::AUR1-C*

*NDT80::hphNT1::P4×lexA-9×Myc-NDT80* /*NDT80::hphNT1::P4×lexA-9×Myc-NDT80*	^1^

YSIY793	*MAT***a**/*MAT*α *his3*Δ*SK*/*his3*Δ*SK ura3*/*ura3::pRS306-mCherry-Spo20^51-91^ trp1::hisG*/*trp1::hisG leu2*/*leu2 arg4-NspI/ARG4*

*lys2/lys2 ho*Δ*::LYS2/ho*Δ*::LYS2 rme1::LEU2/RME1*

*AUR1::P_ACT1_-LexA-ER-haVP16::AUR1-C/AUR1::P_ACT1_-LexA-ER-haVP16::AUR1-C*

*NDT80::hphNT1::P4×lexA-9×Myc-NDT80* /*NDT80::hphNT1::P4×lexA-9×Myc-NDT80* *gip1*∆*::kanMX6/gip1*∆*::kanMX6*	^1^

Plasmid Name	Description	Source

pRS306-mCherry-Spo20^51-91^	*URA3, integration, P_TEF1_-mCherry-Spo20^51-91^*	^1^

1.	Suda, Y., Tachikawa, H., Suda, T., Kurokawa, K., Nakano, A., and Irie, K. (2024). Remodeling of the secretory pathway is coordinated with de novo membrane formation in budding yeast gametogenesis. iScience *27*, 110855. https://doi.org/10.1016/j.isci.2024.110855.
